## supporting information for "AI-enabled Alkaline-resistant Evolution of Protein to Apply in Mass Production"

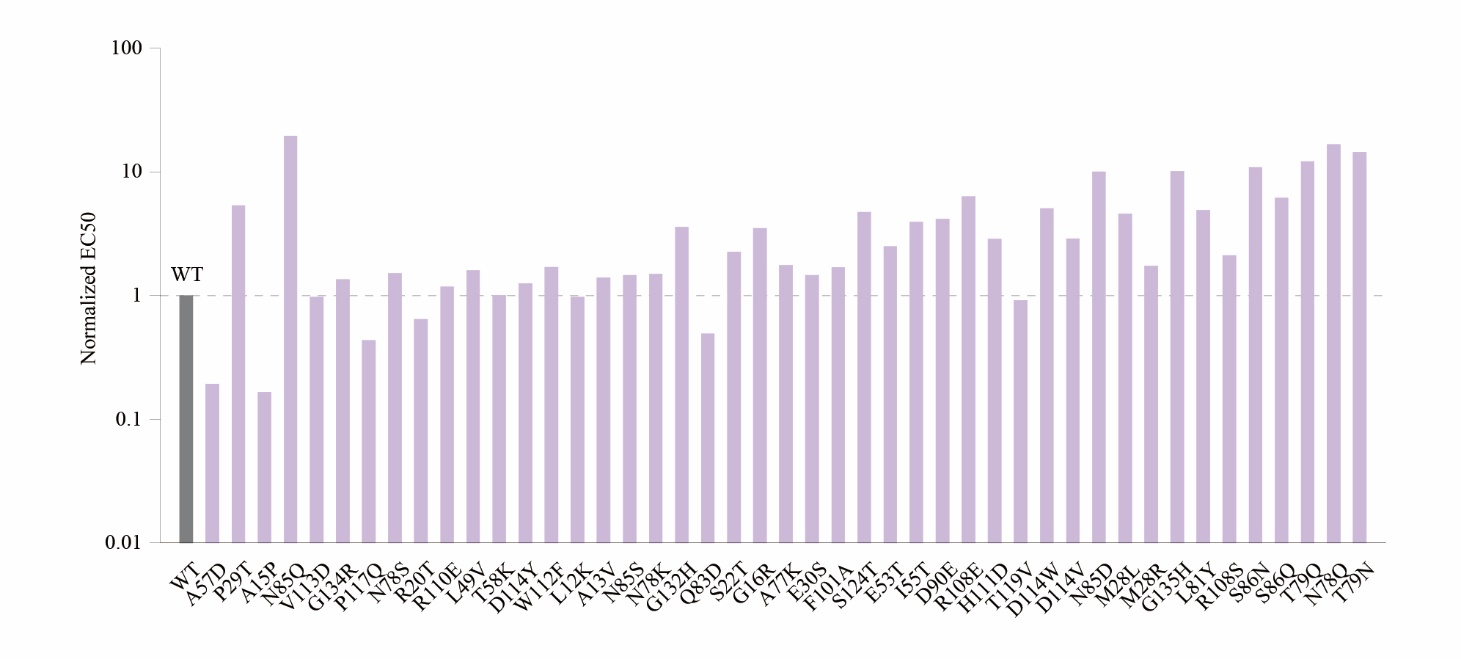

**Figure S1.** Experimental results of single-point mutants before alkali treatment. The affinity of mutants are shown as purple bars. The affinity values were normalized in the graph, with the wild-type EC50 set as 1.

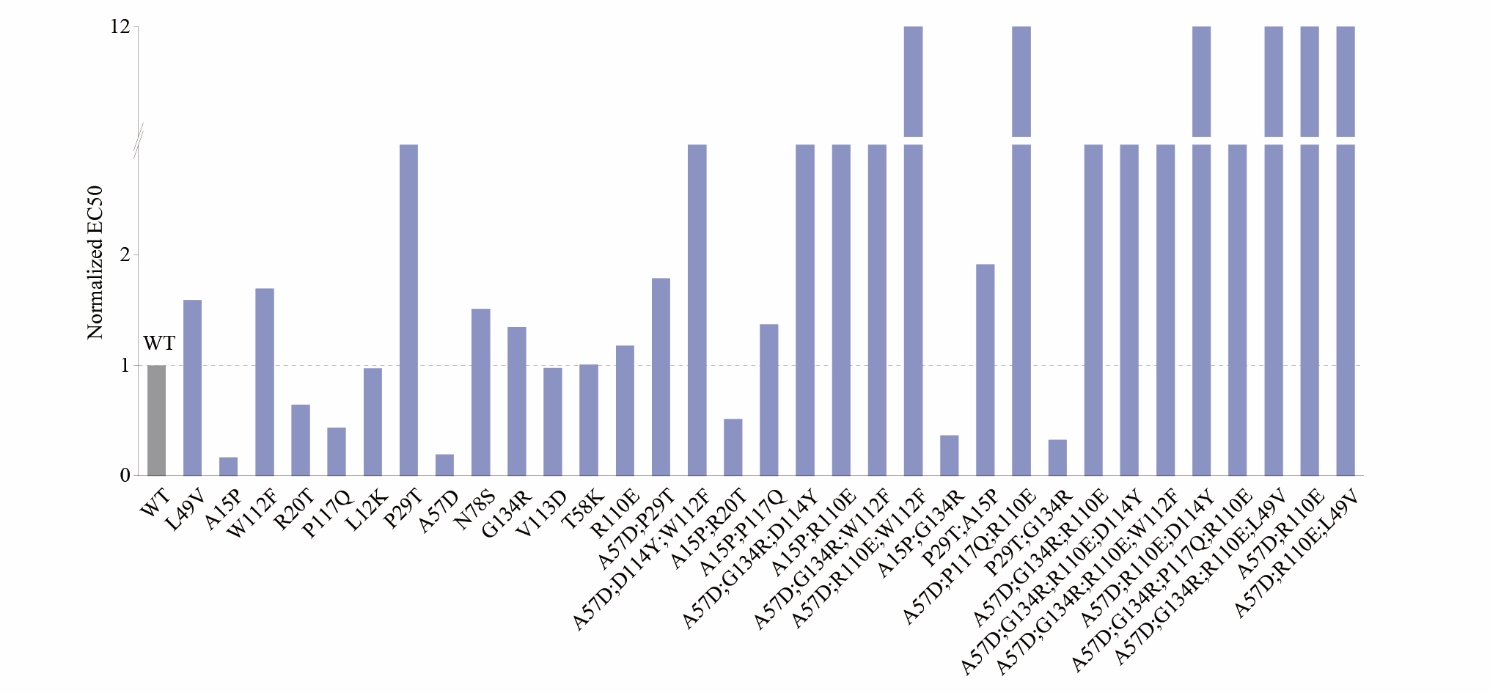

**Figure S2.** Experimental results of multi-point mutants and single-point mutants before alkali treatment. The affinity of mutants are shown as blue bars. The affinity values were normalized in the graph, with the wild-type EC50 set as 1.

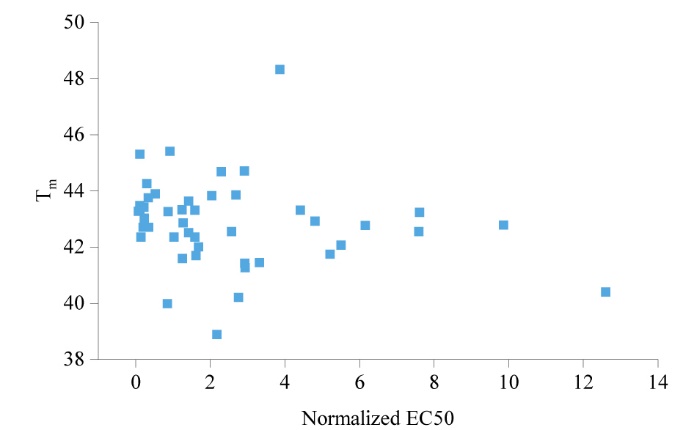

**Figure S3.** Distribution plot of the Normalized EC50 values after 0.3 M NaOH treatment and T_m_ values of single-point mutants. The spearman correlation between the two values is -0.29. The affinity values were normalized in the graph, with the wild-type EC50 set as 1.

**Prediction with Pro-PRIME model**

Pro-PRIME is a deep learning-based methodology developed to guide protein engineering, utilizing a comprehensive dataset comprising 96 million sequence-host bacterial strain optimal growth temperatures (OGT)[1]. Through masked language modeling (MLM) and multi-task learning, Pro-PRIME can learn and comprehend the semantic and grammatical features inherent in protein sequences, and further capture the temperature traits associated with these sequences. As a result, Pro-PRIME naturally correlates higher scores with sequences that are more likely to contribute to robustness and survivability in varied environmental conditions, including extreme temperature scenarios. First, we deployed homologous sequences of the protein as an unsupervised dataset, optimizing both the encoder and masked language modeling modules of Pro-PRIME, since additional unsupervised fine-tuning of the language modeling module on homologous protein sequences of target proteins yields improved results. Then we utilized the Pro-PRIME model to predict zero-shot scores for saturated single-point mutations of VHH antibodies, selecting the top 45 for the first round of experiments. When predicting multi-point mutations, we initially established a library of multi-point mutants based on the results of single-point experiments. We selected 20 single-point mutations for combination into multi-point mutants: A57D, P29T, A15P, N85Q, V113D, G134R, P117Q, N78S, R20T, R110E, L49V, T58K, D114Y, W112F, L12K, A13V, N85S, N78K, G132H, and Q83D, as they exhibited strong alkali resistance in preliminary experiments. Regarding supervised learning, the experimental data of the wild type and all single-point mutations were used as the training set for the Pro-PRIME model. Alkali resistance and thermal stability were used to train two separate Pro-PRIME models, which predicted the alkali resistance scores and thermal stability scores of mutants, respectively. When screening multi-point mutants, we prioritized mutants with higher alkali resistance scores while requiring their thermal stability scores not to be lower than that of the wild type. Although we measured the affinity of proteins in experiments, we did not use this indicator for training the model or screening mutants because excessively high affinity is disadvantageous for purifying growth hormone in practical application.

**Molecular Dynamics (MD) simulations**

The initial structures for molecular dynamics (MD) simulations of both the wild type and the mutant were predicted using AlphaFold3[2]. To simulate experimental conditions, each protein was placed in a cubic water box containing 0.1 M NaCl. The CHARMM27 force field [3] and the TIP4P water model were applied throughout the simulations. After an initial energy minimization of 50,000 steps, the systems were heated and equilibrated for 1 ns in the NVT ensemble at 300 K followed by an additional 1 ns in the NPT ensemble at 1 atm. The production phase then involved 200-ns simulations with periodic boundary conditions, using a 2 fs integration time step. The LINCS algorithm was used to constrain covalent bonds involving hydrogen atoms, while Lennard-Jones interactions were cut off at 10 Å.[4] Electrostatic interactions were computed with the particle mesh Ewald method, using a 10 Å cutoff and a grid spacing of approximately 1.6 Å with a fourth-order spline.[5] Temperature and pressure were regulated by the velocity rescaling thermostat and Parrinello-Rahman algorithm, respectively. [6, 7] All simulations were performed using GROMACS 2020.4 software packages.[8] Both systems have reached equilibrium according to the analyses of root mean squared deviation (RMSD).

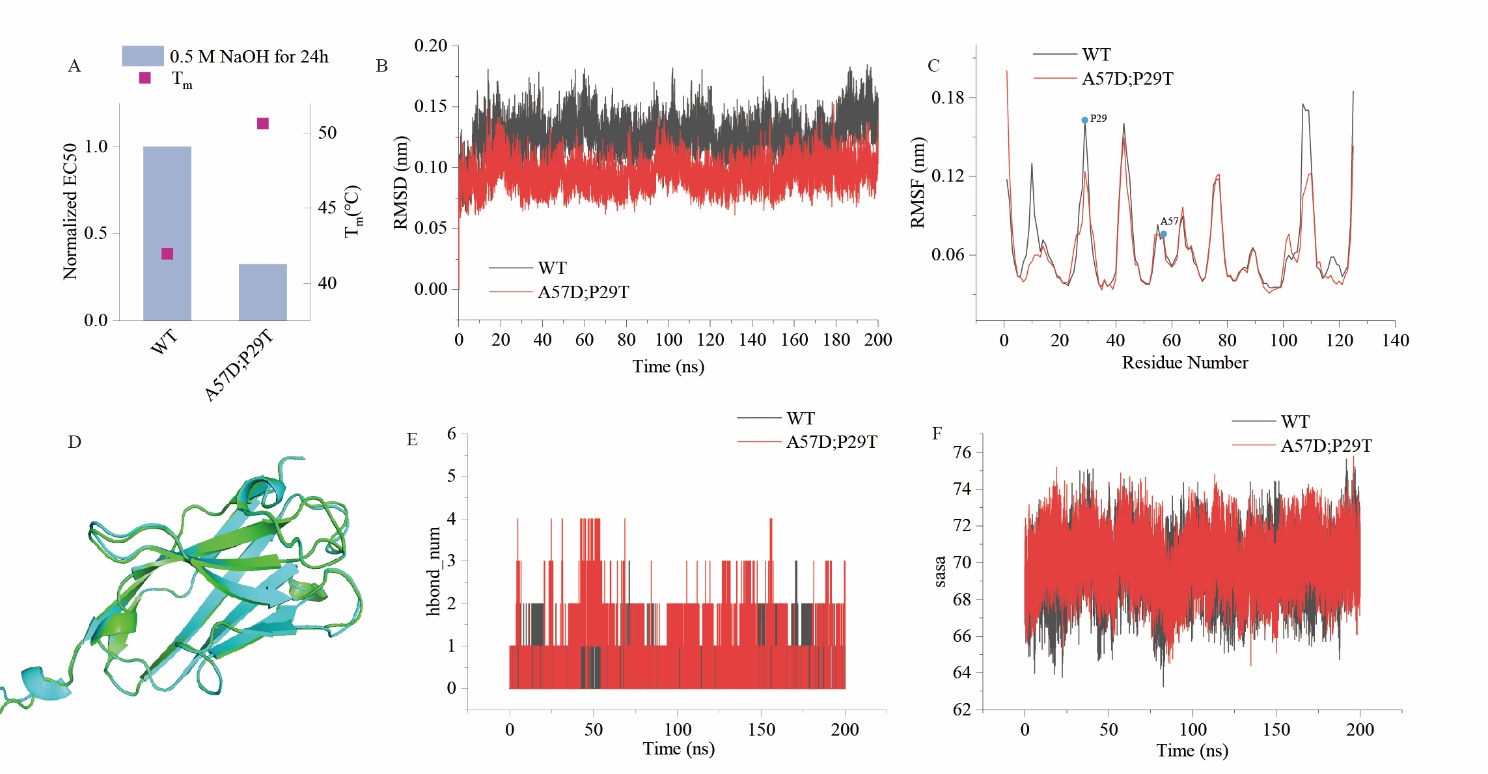

**Figure S4.** **Stability analysis of the wild type (WT) and the A57D;P29T mutant based on MD simulations.** (A) The melting temperatures and binding affinity of the WT and the A57D;P29T mutant after treatment with 0.5 M NaOH for 24 hours. (B) Root-mean-squared deviation (RMSD) for 200ns in MD simulations for both the WT and the mutant. (C) Root-mean-square fluctuations (RMSF) for each residue in the WT and the mutant. (D) Comparison of the AlphaFold3 predicted structures of the WT and the mutant. (E) Comparison of hydrogen bonds and (F) solvent accessible surface area(SASA) in the WT and the mutant.

**Plasmid construction and Protein expression**

A codon-optimized version of VHH antibody and variants genes were synthesized by Sangon Biotech (Shanghai, China). It was cloned into the pET29(a) plasmid to construct pET29a-VHH-MX with an N terminal His-tag. The expression plasmid was transformed into *E.coli* BL21(DE23) cells. A single colony of each recombinant *E. coli* strain was inoculated into 30 mL LB medium with 50 μg/mL kanamycin for seed culture at 37 °C for 12-16 h. The seed culture was transferred 10 ml to 1L LB medium with 50 µg/ml kanamycin at 37 °C 220 rpm until the OD_600_ value reached 0.6-0.8. The culture was cooled to 16 °C and then induced with 0.5 mM IPTG for 20-24 h at 16 °C.

**Protein Purification**

Cells were harvested from the fermentation culture by centrifugation for 30 min at 4,000 rpm, and the cell pellets were collected for later purification. The cell pellets were resuspended in buffer A (20 mM Na_2_HPO_4_ and NaH_2_PO_4_, 0.5 M NaCl, pH 8.0) and lysed using ultrasonication. The lysates were centrifuged at 12,000 rpm at 4 °C for 30 min, after which the supernatants were subjected to Ni-NTA affinity purification with elution buffer (20 mM Na_2_HPO_4_ and NaH_2_PO_4_, 0.5 M NaCl, 250 mM imidazole, pH 8.0). The purity of the fractions obtained was analyzed using SDS-PAGE. The fractions containing the purified target protein were combined and desalted using an ultrafiltration unit. The concentrated protein was finally stored in buffer A supplemented with 10% glycerol at -80°C for long-term preservation.

**Affinity test (ELISA)**

Ninety-six-well plates were coated with growth hormone protein at a density of 5 ng/well at 4 °C overnight. The plates were washed with 1 × PBS’T three times. Following blocking with 1% BSA in 1 × PBS at 25°C for 2 h. After washing three times with 1 × PBS’T, the plates were incubated with serial dilutions of VHH proteins 100 μl/well (1:2,1:4, 1:8, 1:16, 1:32, 1:64, 1:128, 1:256, 1:512, 1:1024, 1:2048) for 1 h at 25°C. After washing three times with 1 × PBS’T, 100 μl/well HRP-labeled Goat Anti-Mouse IgG(H+L) (1:5000) were added and incubated at 25°C for 1h. The plates were washed with 1 × PBS’T four times, a total of 100 μl/well 3,3′,5,5′-tetramethylbenzidine was added and incubated at 25 °C for 15 min in the dark. Finally, 100 μl/well 2 M H_2_SO_4_ was added to stop the reaction and absorbance was measured at 450 nm (TECAN, Swiss).

The log(agonist) vs. response -- Variable slope (four parameters) curves were analyzed to calculate EC50 which determines the stability of VHH after alkaline treatment.

**Differential scanning fluorimetry (DSF)**

The thermal stability testing was carried out using a PCR instrument (Analytik Jena qTower3). All proteins were diluted in 1× PBS to a final concentration of 0.3 mg/ml and mixed with SYPRO Orange at a final concentration of 5× in an eight-row PCR tube. The protein unfolding process was initiated by subjecting the samples to a thermal treatment ranging from 25 to 85 °C (with a temperature increment of 0.5 °C/step) with each step holding for 5 seconds. Subsequently, the thermal unfolding curves were obtained, and the data were analyzed using the Boltzmann equation to determine the melting temperature (T_m_).

**Alkaline pH stability test (SDS-PAGE)**

The VHH antibodies (0.5 mg/mL) were treated an equal volume of 0.5 M NaOH for 24 hours, followed by pH adjustment using the same volume of 0.5 M HCl. After centrifugation, the supernatant was collected and analyzed by SDS-PAGE, and the gray intensity of the fragmented bands was analyzed using ImageJ to determine the alkaline cleavage status of the VHH antibodies.

**Alkaline pH stability test (ELISA)**

The VHH antibodies (1.5 mg/mL) were treated an equal volume of 0.3 M or 0.5 M NaOH for 24 hours, followed by pH adjustment using the same volume of 0.5 M HCl, the supernatant was collected after centrifugation.

Ninety-six-well plates were coated with growth hormone protein at a density of 5 ng/well at 4 °C overnight. The plates were washed with 1 × PBS’T three times. Following blocking with 1% BSA in 1 × PBS at 25°C for 2 h. After washing three times with 1 × PBS’T, the plates were incubated with serial dilutions of VHH proteins 100 μl/well (1:2,1:4, 1:8, 1:16, 1:32, 1:64, 1:128, 1:256, 1:512, 1:1024, 1:2048) for 1 h at 25°C. After washing three times with 1 × PBS’T, 100 μl/well HRP-labeled Goat Anti-Mouse IgG(H+L) (1:5000) were added and incubated at 25°C for 1h. The plates were washed with 1 × PBS’T four times, a total of 100 μl/well 3,3′,5,5′-tetramethylbenzidine was added and incubated at 25 °C for 15 min in the dark. Finally, 100 μl/well 2 M H_2_SO_4_ was added to stop the reaction and absorbance was measured at 450 nm (TECAN, Swiss).

The log(agonist) vs. response -- Variable slope (four parameters) curves were analyzed to calculate EC50 which determines the stability of VHH after alkaline treatment.

**Acid Dissociation test (ELISA)**

Ninety-six-well plates were coated with growth hormone protein at a density of 5 ng/well at 4 °C overnight. The plates were washed with 1 × PBS’T three times. Following blocking with 1% BSA in 1 × PBS at 25°C for 2 h. After washing three times with 1 × PBS’T, the plates were incubated with serial dilutions of VHH proteins 100 μl/well (1:2,1:4, 1:8, 1:16, 1:32, 1:64, 1:128, 1:256, 1:512, 1:1024, 1:2048) for 1 h at 25°C. After washing three times with 20 mM Citric Acid and one time with 1 × PBS’T, 100 μl/well HRP-labeled Goat Anti-Mouse IgG(H+L) (1:5000) were added and incubated at 25°C for 1h. The plates were washed with 1 × PBS’T four times, a total of 100 μl/well 3,3′,5,5′-tetramethylbenzidine was added and incubated at 25 °C for 15 min in the dark. Finally, 100 μl/well 2 M H_2_SO_4_ was added to stop the reaction and absorbance was measured at 450 nm (TECAN, Swiss).

The log(agonist) vs. response -- Variable slope (four parameters) curves were analyzed to calculate EC50 which determines the stability of VHH after acid dissociation.

**Salt Dissociation test (ELISA)**

Ninety-six-well plates were coated with growth hormone protein at a density of 5 ng/well at 4 °C overnight. The plates were washed with 1 × PBS’T three times. Following blocking with 1% BSA in 1 × PBS at 25°C for 2 h. After washing three times 1 × PBS’T, the plates were incubated with serial dilutions of VHH proteins 100 μl/well (1:2,1:4, 1:8, 1:16, 1:32, 1:64, 1:128, 1:256, 1:512, 1:1024, 1:2048) for 1 h at 25°C. After washing three times with 20mM PB buffer (20 mM Na_2_HPO_4_ and NaH_2_PO_4_, 1 M NaCl, pH 8.0) and one time with 1 × PBS’T, 100 μl/well HRP-labeled Goat Anti-Mouse IgG(H+L) (1:5000) were added and incubated at 25°C for 1h. The plates were washed with 1 × PBS’T four times, a total of 100 μl/well 3,3′,5,5′-tetramethylbenzidine was added and incubated at 25 °C for 15 min in the dark. Finally, 100 μl/well 2 M H_2_SO_4_ was added to stop the reaction and absorbance was measured at 450 nm (TECAN, Swiss).

The log(agonist) vs. response -- Variable slope (four parameters) curves were analyzed to calculate EC50 which determines the stability of VHH after salt dissociation.

**Acid Resistance test (ELISA)**

The VHH antibodies (1.5 mg/mL) were treated an equal volume of 1M ethanoic acid for 48 hours, followed by pH adjustment using the same volume of 1 M NaOH, the supernatant was collected after centrifugation.

Ninety-six-well plates were coated with growth hormone protein at a density of 5 ng/well at 4 °C overnight. The plates were washed with 1 × PBS’T three times. Following blocking with 1% BSA in 1 × PBS at 25°C for 2 h. After washing three times with 1 × PBS’T, the plates were incubated with serial dilutions of VHH proteins 100 μl/well (1:2,1:4, 1:8, 1:16, 1:32, 1:64, 1:128, 1:256, 1:512, 1:1024, 1:2048) for 1 h at 25°C. After washing three times with 1 × PBS’T, 100 μl/well HRP-labeled Goat Anti-Mouse IgG(H+L) (1:5000) were added and incubated at 25°C for 1h. The plates were washed with 1 × PBS’T four times, a total of 100 μl/well 3,3′,5,5′-tetramethylbenzidine was added and incubated at 25 °C for 15 min in the dark. Finally, 100 μl/well 2 M H_2_SO_4_ was added to stop the reaction and absorbance was measured at 450 nm (TECAN, Swiss).

The log(agonist) vs. response -- Variable slope (four parameters) curves were analyzed to calculate EC50, which determines the stability of VHH after acid treatment.

**Alkaline incubation of VHH followed by Liquid Chromatography Mass Spectrometry (LC-MS) analysis**

The VHH antibodies (1.5 mg/mL) were treated an equal volume of 0.5 M NaOH for 24 hours, followed by pH adjustment using the same volume of 0.5 M HCl, the supernatant was collected for LC-MS after centrifugation. The LC-MS analyses were performed using a Waters Xevo G2 Q-TOF MS instrument (Waters, Milford, MA, US) equipped with an AQUITY H-class UPLC system (Waters). The MS instrument was calibrated daily with sodium iodide standard provided by the vendor (2 µg/µL in 50:50 2-Propanol: Water, 700001593). All data was collected using a Zorbax 300SB-C8 (2.1 × 50 mm 1.7 µm) column (Agilent Technologies, Santa Clara, CA, US). The column was operated at 60 °C and 5 µL of each sample was injected (approximately 100–150 nmol on column). The LC gradient consisted of Buffer B (0.1% formic acid in water) and Buffer C (0.1% formic acid in 80:20 Acetonitrile:2-propanol). The gradient was 0–2 min 100% B, 2–10 min 100–10% B, 10–11 min 0% B, 11–12 min 100% B at a flow rate of 0–11 min 0.5 mL/min and 11–12 min 1 mL/min. The MS was run in positive polarity and Sensitivity mode at a m/z range of 50–2000, scan rate of 1 spectrum/second and ESI potential set at 3 kV. Determination of molecular weight (MW) for each peak in the chromatograms was performed by the provided MaxEnt1 deconvolution function in the MassLynx software (Waters).

Reference:

[1] F. Jiang, M. Li, J. Dong, Y. Yu, X. Sun, B. Wu, J. Huang, L. Kang, Y. Pei, L. Zhang, A general temperature-guided language model to design proteins of enhanced stability and activity, Science Advances 10(48) (2024) eadr2641.

[2] J. Abramson, J. Adler, J. Dunger, R. Evans, T. Green, A. Pritzel, O. Ronneberger, L. Willmore, A.J. Ballard, J. Bambrick, Accurate structure prediction of biomolecular interactions with AlphaFold 3, Nature (2024) 1-3.

[3] P. Bjelkmar, P. Larsson, M.A. Cuendet, B. Hess, E. Lindahl, Implementation of the CHARMM force field in GROMACS: analysis of protein stability effects from correction maps, virtual interaction sites, and water models, J. Chem. Theory Comput. 6(2) (2010) 459-466.

[4] B. Hess, C. Kutzner, D. Van Der Spoel, E. Lindahl, GROMACS 4: algorithms for highly efficient, load-balanced, and scalable molecular simulation, J. Chem. Theory Comput. 4(3) (2008) 435-447.

[5] U. Essmann, L. Perera, M.L. Berkowitz, T. Darden, H. Lee, L.G. Pedersen, A smooth particle mesh Ewald method, J. Chem. Phys. 103(19) (1995) 8577-8593.

[6] G. Bussi, D. Donadio, M. Parrinello, Canonical sampling through velocity rescaling, J. Chem. Phys. 126(1) (2007).

[7] M. Parrinello, A. Rahman, Polymorphic transitions in single crystals: A new molecular dynamics method, J. Appl. Phys. 52(12) (1981) 7182-7190.

[8] M.J. Abraham, T. Murtola, R. Schulz, S. Páll, J.C. Smith, B. Hess, E. Lindahl, GROMACS: High performance molecular simulations through multi-level parallelism from laptops to supercomputers, SoftwareX 1 (2015) 19-25.
